# Cryo-EM reveals a mechanism of USP1 inhibition through a cryptic binding site

**DOI:** 10.1101/2022.04.06.487267

**Authors:** ML Rennie, C Arkinson, V Chaugule, H Walden

## Abstract

Repair of DNA damage is critical to genomic integrity and frequently disrupted in cancers. USP1, a nucleus-localized deubiquitinase, lies at the interface of multiple DNA repair pathways and is a promising drug target for certain cancers. Although multiple inhibitors of this enzyme, including one in phase I clinical trials, have been established, their binding mode is unknown. Here we use cryo-Electron Microscopy to study an assembled enzyme-substrate-inhibitor complex of USP1 and the well-established inhibitor, ML323. Achieving 2.5 Å resolution, we discover an unusual binding mode in which the inhibitor displaces part of the hydrophobic core of USP1. The consequent conformational changes in the secondary structure lead to subtle rearrangements in the active site that underlie the mechanism of inhibition. These structures provide a platform for structure-based drug design targeting USP1.

**One Sentence Summary:** USP1, a cancer target, is inhibited by ML323 displacing part of the protein fold, allosterically disrupting the active site.

## Main Text

Human DNA is under constant assault from external and endogenous agents that induce damage, including ultraviolet (UV) radiation, smoking, replication stress, cellular metabolites, and the radiation and chemotherapeutics widely used in targeted cancer treatments. In response, multiple organized DNA repair pathways have evolved, with different types of DNA damage eliciting different responses. Ubiquitin-specific protease 1 (USP1) is a nucleus-localized deubiquitinase and a well-established component of DNA repair, acting in the Fanconi Anemia (FA) pathway (*1*) and translesion synthesis (TLS) (*2*) to catalyze the removal of specific monoubiquitin signals (Fig. 1A). In the FA pathway USP1 deubiquitinates the FANCI-FANCD2 heterodimer to release the complex from DNA (*1*, *3*), while in TLS it acts on Proliferating Cell Nuclear Antigen (PCNA) to regulate polymerase recruitment (*2*). USP1 has also been shown to deubiquitinate several oncogenic proteins to prevent their degradation (*4, 5*).

**Fig. 1.**
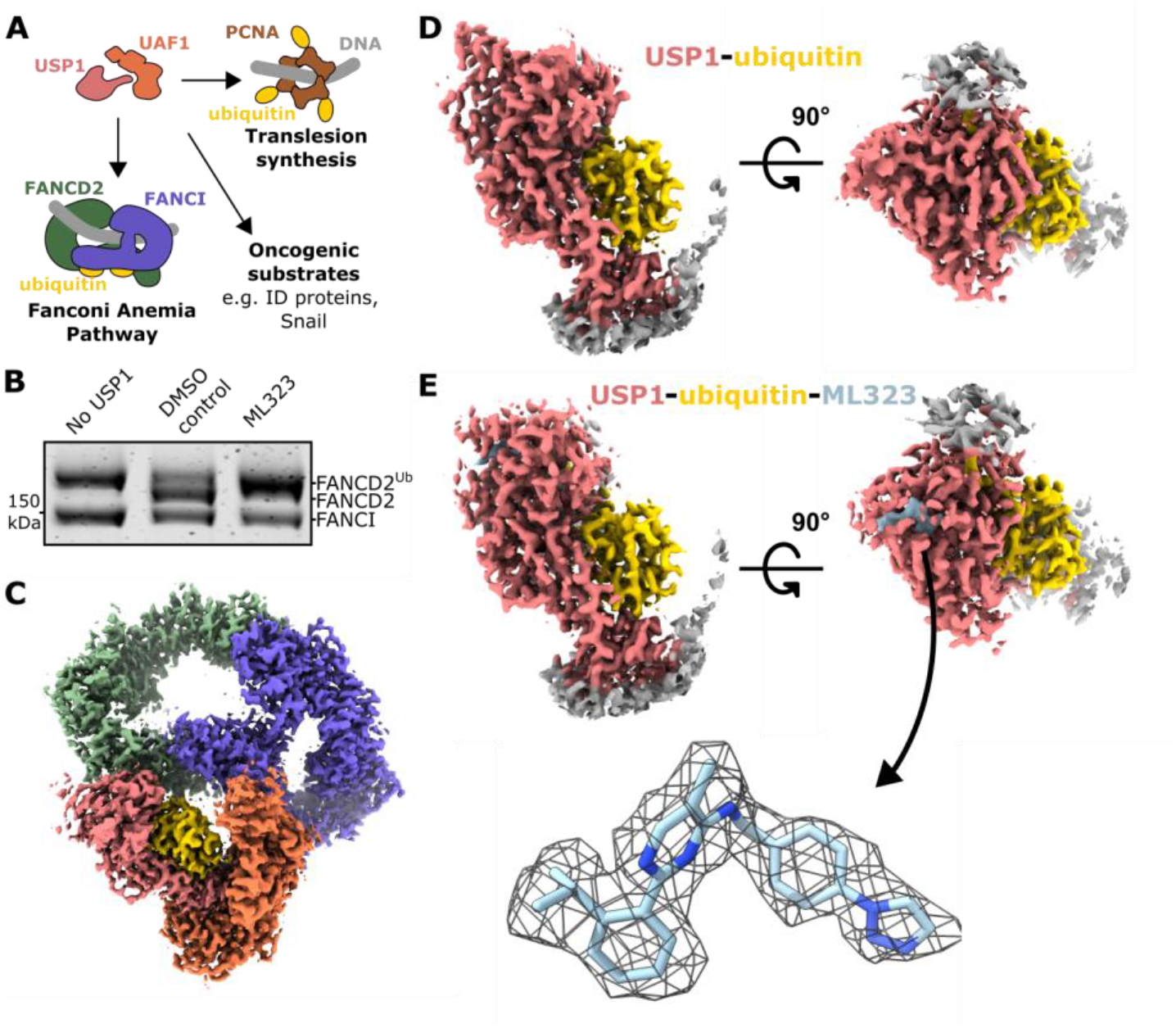
Cryo-EM structures of inhibitor-bound and unbound USP1. (**A**) Schematic representation of USP1-UAF1 and its substrates. (**B**) ML323 inhibits deubiquitination of FANCI-FANCD2^Ub^ by USP1-UAF1 *in vitro*. Reactions of USP1-UAF1 at 0.1 μM and FANCI-FANCD2^Ub^ at 1 μM in the presence or absence of 10 μM ML323 (0.25% DMSO) were terminated at 10 min and assessed by SDS-PAGE and Coomassie staining. Image is representative of two replicates. (**C**) Cryo-EM map of reconstituted FANCI (purple), FANCD2 (green) conjugated with ubiquitin (yellow), USP1 (pink), UAF1 (orange), and ML323. (**D**) Focused map of USP1-ubiquitin without ML323. (**E**) Focused map of USP1-ubiquitin with ML323 (blue). A refined model of ML323 and surrounding map contoured at 6.8 rmsd is highlighted.

USP1 is a promising target for the treatment of several types of cancer. USP1 expression is upregulated in certain breast cancers (*6, 7*), ovarian cancers (*5*), colorectal cancers (*8*), and bone cancers (*4*), often correlated with poor prognosis (*5*, *7*, *8*). Reduction of USP1 activity via small molecules or at the level of gene expression has been shown to decrease growth of cancer cells (*4*), particularly when in combination with other DNA-targeting treatments (*5*, *6, 8–10*). This has fueled interest in development of USP1 inhibitors (*9–11*), with the first phase I clinical trials commencing in 2021 (*12*). However, the mechanism for inhibition remains unknown due to a lack of structural data. In particular there are no available enzyme-inhibitor structures. ML323, a well-established inhibitor, is thought to interact with USP1 allosterically (*10*) and, at the cellular level, has been suggested to cause replication stress via trapping of USP1 on DNA (*13*). Identification of the proposed allosteric binding site has remained elusive hampering efforts to develop the next generation of inhibitors.

Recently, we determined the first structures of USP1 using crystallography and cryo-EM at resolutions sufficient to resolve protein sidechains (3.2-3.7 Å) (*14*). Although such resolution provides understanding of protein-protein interfaces, it is not high enough to model features required in structure-guided drug design, which typically needs 2.5 Å or better to unambiguously identify small molecule ligands and their interactions (*15*). Despite extensive efforts, we have found it challenging to produce crystals of USP1 diffracting to high resolution, likely due to inherent flexibility. The small size of USP1 (90 kDa) also makes it a difficult target for cryo-EM. However, when assembled with both its activating partner UAF1 (*16*), and its largest substrate, the FANCI-FANCD2 DNA-clamp (*17*–*20*), the complex reaches 0.5 MDa. We recently reported the structure of this complex, mutated at the active site cysteine to limit catalysis (Cys90Ser) (*14*). Although still at limited global resolution, we were able to resolve the active site, including the isopeptide bond of the ubiquitinated substrate, so we wondered if this scaffold would be amenable to identifying the binding site and mechanism of USP1 inhibition by ML323. Previous studies have focused on di-ubiquitin and ubiquitinated PCNA as substrates in inhibitor screens, therefore we first asked whether ML323 inhibits FANCI-FANCD2 deubiquitination *in vitro*. We find that, in our reconstituted assay, the FANCI-FANCD2 heterodimer, with monoubiquitin on FANCD2, is readily deubiquitinated by USP1-UAF1, but this is significantly reduced in the presence of ML323 (Fig. 1B). We then assembled the enzyme-substrate-inhibitor complex from the 6 individual components (USP1^C90S^-UAF1-FANCI-FANCD2^Ub^-dsDNA-ML323). Single particle analysis of this complex with cryo-EM yields a consensus reconstruction to 2.7 Å (Fig 1C, fig. S1, table S1). Focused refinement of USP1-ubiquitin and classification into two states yields reconstructions of USP1 with and without ML323, each to 2.5 Å (Fig. 1D-E, fig. S2). At this resolution the inhibitor can be unambiguously identified and modeled as well as several water molecules within the structures (Fig. 1E, fig. S3).

Rather than binding to a pocket on the surface of USP1, or in the active site, the inhibitor binds within the hydrophobic core of USP1 by displacing and replacing elements of the USP-fold (Fig. 2A-B). More than 80% of the ML323 binding site is inaccessible to solvent in the unbound state (Fig 2C). This cryptic site lies between the palm and thumb subdomains of the USP-fold, ~9 Å from the active site cysteine (Cys90 mutated to Ser; Cβ to nearest non-hydrogen atom of ML323). ML323 displaces two short β-strands of the thumb, contacting numerous hydrophobic residues of both the palm and thumb that were previously part of the hydrophobic core (Fig. 2B,D). One of the β-strands resides at the N-terminus of the USP fold and is completely displaced, the other becomes less ordered (Fig. 2D). In addition, the sidechain of Gln97 is displaced by ML323 to form a hydrogen bond with a water molecule and may form a strained hydrogen bond with the pyrimidine of ML323. The isopropyl group of ML323 occupies the same position of the equivalent group of Leu83 in the unbound structure, while the triazole contributes a hydrogen bond with Gln160, explaining why these modifications improved inhibition compared to the initial inhibitor on which ML323 was based (*10*, *11*) (Fig. 2B, fig. S4). The helix on which Gln160 is located, bends to form additional van der Waals contacts with ML323. This helix and the adjoining loop is shorter in two closely related USPs which also interact with UAF1, USP12 (*21*, *22*) and USP46 (*23*) (Fig. 2E), as well as in other USPs (fig. S5). In effect, the inhibitor replaces a buried segment of the protein structure and increases disorder in the surrounding region.

**Fig. 2.**
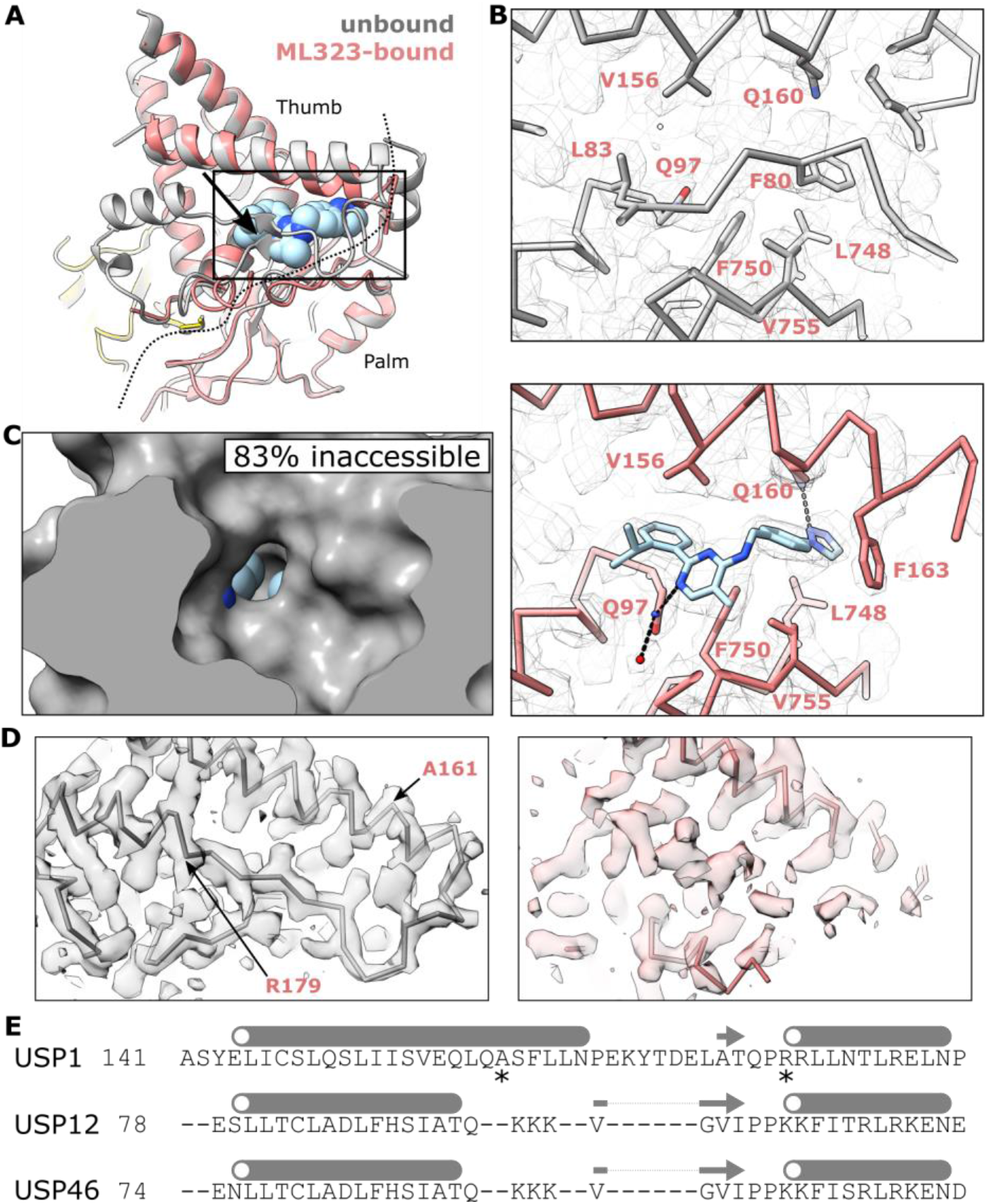
Inhibitor binding displaces the hydrophobic packing of USP1. (**A**) Superposition of the ML323-bound (colored) and unbound (gray) structures. A dashed line represents the interface between the palm and thumb. The displaced β-strands of the thumb are indicated with an arrow. (**B**) Structural detail of the ML323 binding site. Q160 forms a potential hydrogen bond with ML323, shown as a dashed line. Cα trace and side chains involved in ML323 binding are shown. Cryo-EM maps are shown as mesh contoured at 6.8 rmsd. (**C**) Solvent-excluded surface of USP1 from the unbound structure with ML323 (spheres) from the bound structure overlaid. The percentage of ML323 buried within the solvent-excluded surface is indicated. (**D**) Region of USP1 that becomes less ordered upon ML323 binding. Cα trace is shown and cryo-EM maps as transparent solid contoured at 6.8 rmsd. (**E**) Structure-based sequence alignment of USP1 with close homologs. Region in (**D**) is shown with highlighted residues indicated with an asterisk.

The structures also explain the mechanism of inhibition by ML323. Although the active site cysteine is not significantly perturbed in the ML323-bound structure, there is a subtle change in the loop containing the catalytic aspartate, Asp751, on the palm subdomain (Fig. 3A). This disruption breaks the hydrogen bond of the catalytic aspartate with the catalytic histidine, His593, and instead an adjacent aspartate, Asp752, forms a hydrogen bond with the histidine causing it to flip and form an additional hydrogen bond with a water molecule (Fig. 3B). In the unbound state this histidine is poised to deprotonate what would be the catalytic cysteine which would then allow nucleophilic attack on the isopeptide (Fig. 3B). However, in the flipped conformation, when ML323 is bound, the histidine is unable to deprotonate the cysteine. This flipping likely underlies the reduction in the catalytic rate by ML323 (Fig. 1B). Reduced binding of ML323 to substrate-bound USP1 that has been observed (*10*) may arise due the new position of the displaced β-strands segment introducing steric hinderance with the tail of ubiquitin and adjoining lysine or the substrate itself decreasing the stability of the enzyme-substrate-inhibitor complex (fig. S6). ML323 can therefore be classified as a Type IV deubiquitinase inhibitor (*24*) – binding outside of the ubiquitin site and allosterically inhibiting catalysis. Overall, the structures provide an allosteric mechanism that rationalizes the mixed mode of inhibition.

**Fig. 3.**
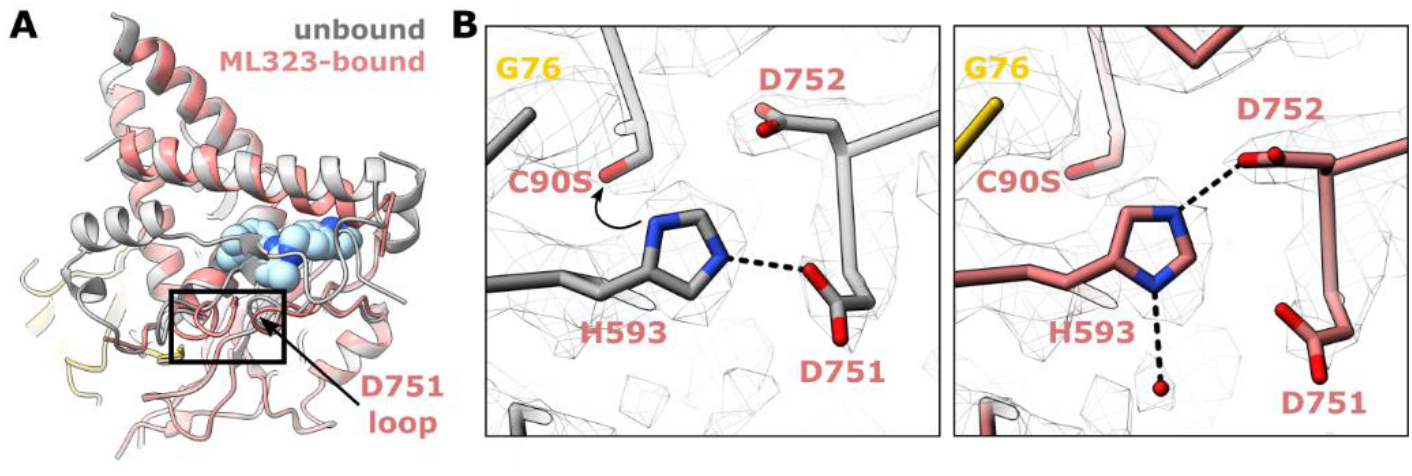
Inhibitor binding disrupts the catalytic site. (**A**) The loop on which the catalytic aspartate, D751, is pushed out by ML323. (**B**) The catalytic histidine, H593, and cysteine, C90S, remain in approximately the same position, however D752 replaces the hydrogen bond with H593 causing it to flip. Hydrogen bonds are shown as dashed lines. Nucleophilic attack is shown by a solid arrow.

USP1 contains multiple insertions in the USP core fold (Fig. 4A). While these are predicted to be structurally disordered and are not tightly associated with the USP fold (*14*), they play important roles in regulating USP1 function (*2*, *13*, *25*, *26*). The N-terminal extension (NTE; residues 1-75) of USP1 has been shown to discriminate between FANCD2 and other substrates by interacting with and facilitating deubiquitination of FANCD2 (*25*). However, the structural basis of this discrimination is unknown. We used AlphaFold-multimer (*27*, *28*) to generate a model of USP1^NTE^ with FANCD2 (Fig. 4B, fig. S7). A short segment of USP1^NTE^, residues 21-25, exhibited good confidence scores (pLDDT>70) (Fig. 4B-C). This region maps well to previous biochemical experiments, in which mutation of these residues to alanine disrupted FANCD2 deubiquitination (*25*). Focused refinement and LocSpiral postprocessing (*29*) of the USP1-UAF1-FANCD2^Ub^-FANCI-dsDNA cryo-EM data around this region showed additional density in the same region as the NTE of the AlphaFold model (Fig. 4B, fig. S8). In the AlphaFold model, Leu23 of the NTE inserts into a hydrophobic pocket of FANCD2, with Arg22 forming a salt bridge with Asp667 on FANCD2. The equivalent region in the FANCD2 paralog, FANCI, is more tightly packed (fig. S9). This explains how the NTE of USP1 acts as an allosteric adaptor to target FANCD2 over FANCI.

**Fig. 4.**
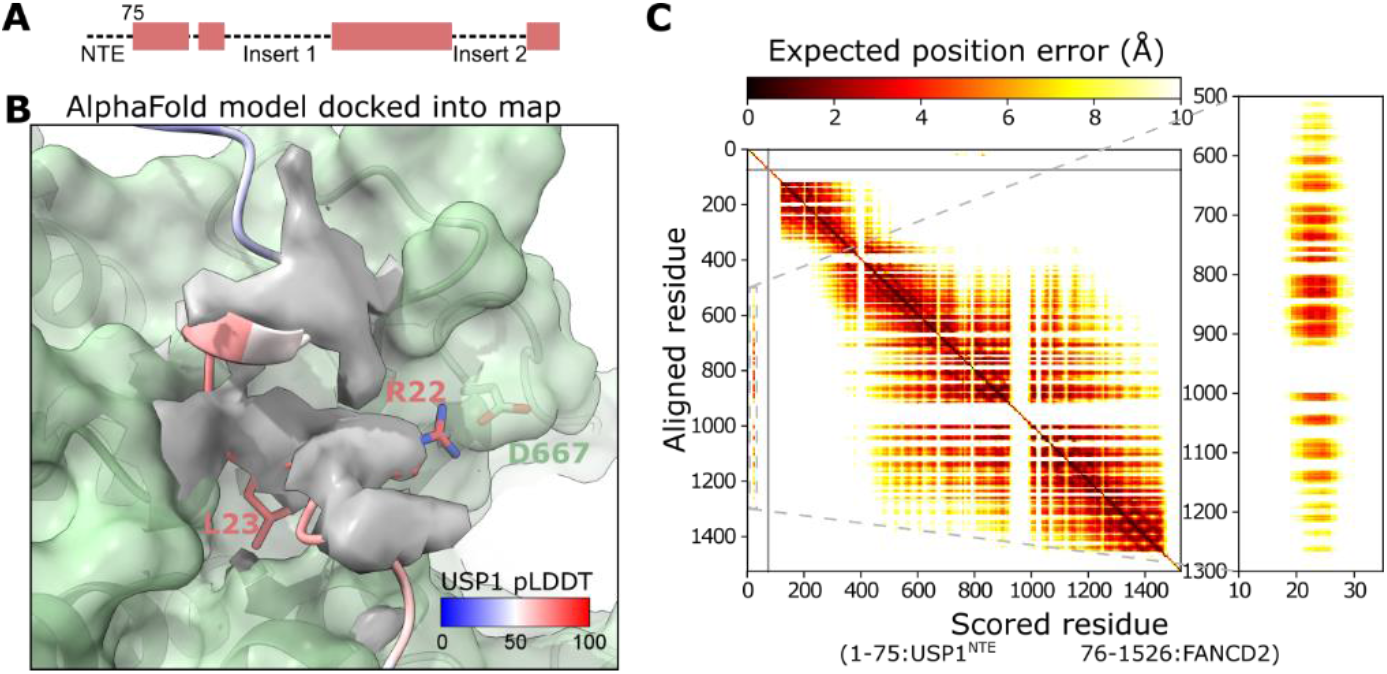
AlphaFold confidently predicts an interaction between USP1^NTE^ and FANCD2. (**A**) Schematic representation of the USP1 primary structure. (**B**) Rigid body fit of an AlphaFold-multimer (*27*) model of USP1^NTE^ and FANCD2 into the cryo-EM map with focused refinement on FANCD2 and LocSpiral (*29*) postprocessing. The cryo-EM map around the density unaccounted for by FANCD2 is shown in solid gray at a contour level of 4 rmsd, FANCD2 is shown as green transparent surface and USP1^NTE^ is colored according to pLDDT AlphaFold confidence scores. (**C**) AlphaFold-multimer predicted aligned error for the USP1^NTE^-FANCD2 complex, highlighting the region of USP1 displayed in (**A**). AlphaFold model and predicted aligned error are representative of 5 replicates.

We have revealed an unexpected mode of binding of USP1 by a small molecule inhibitor displacing a substantial portion of the hydrophobic core. At the resolution achieved we define key interactions that rationalize medicinal chemistry optimizations, and conformational changes induced in the catalytic site that underpin inhibition. Given the role of the NTE in regulation, and the allosteric mode of ML323 inhibition, there are exciting opportunities for the next generation of inhibitors of this important enzyme, and in the development of future cancer treatments.

## Acknowledgements

We thank past and current members of the Walden laboratory for experimental suggestions, comments on the manuscript and their support. We acknowledge Diamond Light Source for access and support of the cryo-EM facilities at the UK’s national Electron Bio-imaging Centre (eBIC) [under proposal BI24557], funded by the Wellcome Trust, MRC and BBRSC. We thank Paula da Fonseca and Edward Morris for assistance with collecting the data at eBIC. We acknowledge the Scottish Centre for Macromolecular Imaging (SCMI) and Dr. Mairi Clarke and Dr. James Streetley for assistance with cryo-EM experiments and access to instrumentation, funded by the MRC (MC_PC_17135) and SFC (H17007). We thank Prof. David Bhella for access to computing resources to run AlphaFold. We thank Dr. Rachel Toth for expression plasmids. We thank Mark Meenan, Paul McLaughlin, and Iain Sim for maintenance of the GPU server.

## Funding

European Research Council (ERC-2015-CoG-681582) ICLUb consolidator grant to H.W.

Medical Research Council (MC_UU_120164/12) to H.W.

## Author contributions

M.L.R. and H.W. conceived this work; M.L.R., C.A., V.K.C. purified proteins; M.L.R. performed all experiments and analyzed all data; M.L.R. wrote the manuscript and prepared figures with contributions from H.W. and C.A.; H.W. secured funding and supervised the project.

## Data and materials availability

All constructs are available on request from the corresponding authors. The atomic coordinates and cryo-EM maps have been deposited to the PDB and EMDB and will be released upon publication.

## Supplementary Materials

Materials and Methods

Fig S1 – S9

Table S1 – S2

References (30 – 50)

