## Supplementary Materials for "Cryo-EM reveals a mechanism of USP1 inhibition through a cryptic binding site"

**This PDF file includes:**

**Materials and Methods**

**Figs. S1 to S9**

**Tables S1 to S2**

**Materials and Methods**

Protein expression and purification

Proteins, all human homologs, were prepared as described previously (*14*). Protein purification buffers and columns used are provided in table S2. Briefly, $His_6$-TEV-USP1$^{G670A,G671A}$, $His_6$-TEV-USP1$^{G670A,G671A,C90S}$, $His_6$-3C-UAF1, $His_6$-3C-FANCD2, $His_6$-FANCI, $His_6$-TEV-V5-FANCI were expressed separately in *Sf*21 insect cells. Cells were lysed by sonication, clarified, and purified by Ni-NTA affinity then anion exchange chromatography. At this stage protein aliquots were occasionally flash frozen in liquid nitrogen and stored -80°C. For $His_6$-TEV-USP1$^{G670A,G671A}$ and $His_6$-TEV-USP1$^{G670A,G671A,C90S}$ TEV protease treatment was performed overnight at 1:10 protease to target protein with gentle agitation before subtractive Ni-NTA affinity chromatography. Flow-through was concentrated to ~10 mg/mL and separated by gel filtration. Purified protein was concentrated to 5-15 mg/mL, flash frozen, and stored at -80°C in 10-20 μL single-use aliquots. All steps were performed on ice or at 4°C and completed within 24-36 hours of lysis. FANCD2 was ubiquitinated and purified using an engineered Ube2T and SpyCatcher-SpyTag setup described in detail elsewhere (*30*, *31*). For FANCD2, the $His_6$-3C tag was removed by 3C protease treatment during preparation of the mono-ubiquitinated version.

Protein concentrations were determined using the predicted extinction coefficients at 280 nm (*32*) and absorbance via a Nanodrop. The ratio of 260nm/280nm was ≤0.65 for all protein batches used in subsequent experiments.

Cryo-EM sample preparation

The USP1$^{C90S}$-UAF1-FANCI-FANCD2$^{Ub}$ complex was prepared by mixing the four individually purified subunits 5:5:1:1. The complex was exchanged into EM buffer (20 mM Tris pH 8.0, 150 mM NaCl, 2 mM DTT) using a Bio-Spin P-30 column (Bio-Rad). The concentration of complex was estimated from absorbance at 280 nm and 1.2 equivalents of dsDNA (61 base pairs; TGATCAGAGGTCATTTGAATTCATGGCTTCGAGCTTCATGTAGAGTCGACGGTGCTGGGAT; IDT) per FANCI-FANCD2$^{Ub}$ was added. ML323 was then added at 2 equivalents of USP1-UAF1. Immediately prior to preparing grids the sample was equilibrated to room temperature for 5 min. UltrAuFoil R1.2/1.3 300 mesh grids were glow discharged twice at 45 mA for 60 secs. A 3.5 μL aliquot of 9.3 μM USP1-UAF1 and 1.8 μM FANCI-FANCD2$^{Ub}$ was applied. The grids were blotted for 3.0 secs and vitrified in liquid ethane using a Vitrobot (ThermoFisher) operating at ~95% humidity at 15°C.

Cryo-EM sample data collection and processing

Initial grid screening was performed on a JEM-F200 (JEOL) equipped with a DE-20 detector (Direct Electron) at the Scottish Centre for Macromolecular Imaging (SCMI). For data collection, a Titan Krios (ThermoFisher) located at eBIC (Diamond Light Source) equipped with a K3 detector (Gatan) was used. A total of 10,998 movies were collected using beam-Image shift. All movies were collected in super-resolution mode with 2x binning and a calibrated pixel size of 1.06 Å using EPU (ThermoFisher). Movies were collected with a total dose of ~40 $e^-/Å^2$ over 40 frames at a rate of 15.46 $e^-$/px/sec.

Subsequent processing was performed in Cryosparc v2.13.2 and v3.3 (*33*) (fig. S1-2, S8A). Patch motion correction, patch CTF estimation, and manual curation was performed resulting in 9,344 dose-weighted, motion corrected micrographs. A maximum alignment resolution of 3 Å was used during patch motion correction. Particle picking, *ab initio* model generation, and cleaning to remove junk picks were performed similarly to described previously (*14*). Blob picking was performed on a subset of micrographs with minimum and maximum particle diameters of 150 Å and 250 Å respectively, using an elliptical blob, followed by 2D classification and *ab initio* reconstruction. Templates from 2D classification were then used to pick from all micrographs and 6.7 million images were extracted with a boxsize of 320x320 pixels. Cleaning was performed iteratively and in batches by heterogenous refinement with one good starting model and 2 or 3 “junk” starting models not representing the protein complex of interest, all low pass filtered to 20 Å. Further rounds of heterogenous refinement were performed using the same starting model low-pass filtered at 12 Å, 15 Å, 20 Å, and 30 Å. Between 1 and 8 full passes through the dataset were used in heterogenous refinements. Particles corresponding to the highest resolution class were then re-extracted yielding 1.3 million particles. Another round of heterogenous refinement yielded 1.15 million particles reaching 2.82 Å after non-uniform refinement (*34*). Local motion correction was performed, again with a maximum alignment resolution of 3 Å improving the resolution to 2.76 Å. Global CTF refinement of beam tilt and trefoil (*35*) against the pooled images was performed improving the resolution to 2.72 Å. All Fourier Shell Correlation (FSC) calculations were performed using masks without auto-tightening as the auto-tightened mask tended to exclude some sidechains and water molecules. Local resolutions were calculated using an adaptive window factor of 20.

Local refinement was performed with a mask covering USP1 and ubiquitin using a gaussian priors of 3° over rotation and 2 Å over shifts with marginalization and non-uniform refinement (fig. S2). 3D variability analysis of this locally aligned region (2 modes and 2.8 Å filter resolution) and clustering was used to yield two states – one corresponding ML323 bound and one unbound. 3D classification with two classes was performed using the two cluster models filtered to 8 Å as inputs. The resulting particles were passed through local refinement again.

For the FANCD2 helical and C-terminal domain local refinement was performed with a mask covering this region using a gaussian priors of 3° over rotation and 2 Å over shifts with marginalization and non-uniform refinement (fig. S8A). LocSpiral (*29*), via the COSMIC2 web platform (https://cosmic-cryoem.org/) (*36*) was used to post-process the half-maps using resolutions between 2.47 and 30 Å, a bandwidth of 8, threshold for significant comparison 0.95, and threshold for 3D mask of 0.14.

Modeling building and refinement

ML323-bound and unbound structures were build into the locally refined maps using USP1 and ubiquitin from the previous structure of USP1-UAF1-FANCI-FANCD2$^{Ub}$ (PDB ID: 7AY1) (*14*). Manual model editing was performed using COOT (37). ML323 restraints were calculated using the GRADE web server (http://grade.globalphasing.org/).

Automated refinement against the globally sharpened maps, with the B-factor estimated from the Guinier plot, were performed using phenix real-space refinement (*38*). A refinement resolution of 2.8 Å (FSC=0.5). Bond and angle restraints for the USP1 Zinc finger and were incorporated. Cryo-EM data and model statistics are reported in table S2. The FSC between the model and map was computed using phenix (*39*) (fig. S2). Structures and maps were analysed and figures produced using ChimeraX (*40*). Volume operations and calculations were performed in Blender (https://www.blender.org/) on solvent-excluded surfaces computed in ChimeraX and exported as Wavefront files (.obj). A Boolean modifier was used to generate the difference between the unbound ML323 surface and the unbound USP1 was taken to be the inaccessible volume. Structure-informed multiple sequence alignments were performed using PROMALS3D (*41*).

AlphaFold models were generated using the full-length human FANCD2 and the N-terminal extension of human USP1 (residues 1-75) sequences. AlphaFold v2.1.0 *(27, 28)* with multimer model preset was used to generate five relaxed models.

Deubiquitination assays

Deubiquitination reactions were performed by preparing a 2x substrate mix and a 2x enzyme mix, and mixing these 1:1 to initiate the reaction. Both mixes were setup on ice, and then incubated at room temperature for at least 20 min prior to reaction initiation and during the reaction. The 2x substrate mix was prepared by diluting stocks (≥30 µM) of $FANCD2_{Ub}$, $His_6$-V5-TEV-FANCI, and dsDNA (61 base pairs) with DUB buffer (20 mM Tris pH 8.0, 75 mM NaCl, 5% glycerol, 1 mM DTT). The resulting 2x mix was composed 2 µM $FANCD2_{Ub}$, 2 µM

FANCI, 8 μM dsDNA. The 2x enzyme mixes were prepared by diluting concentrated stocks (≥30 μM) of USP1, $His_6$-3C-UAF1, and ML323 or DMSO control with DUB buffer. The resulting 2x mixes were composed of 200 nM USP1, 200 nM UAF1, 0.5% DMSO, and 20 μM ML323, where included. Aliquots of 4 μL of reaction were terminated at 10 min by addition of 20 μL 1.2x NuPAGE LDS buffer (Thermo Fisher) supplemented with DTT (final concentration 100 mM). SDS-PAGE was performed using Novex 4–12% Tris-glycine gels (Thermo Fisher) and subsequent staining of the gels with Instant- Blue Coomassie stain (Expedeon).

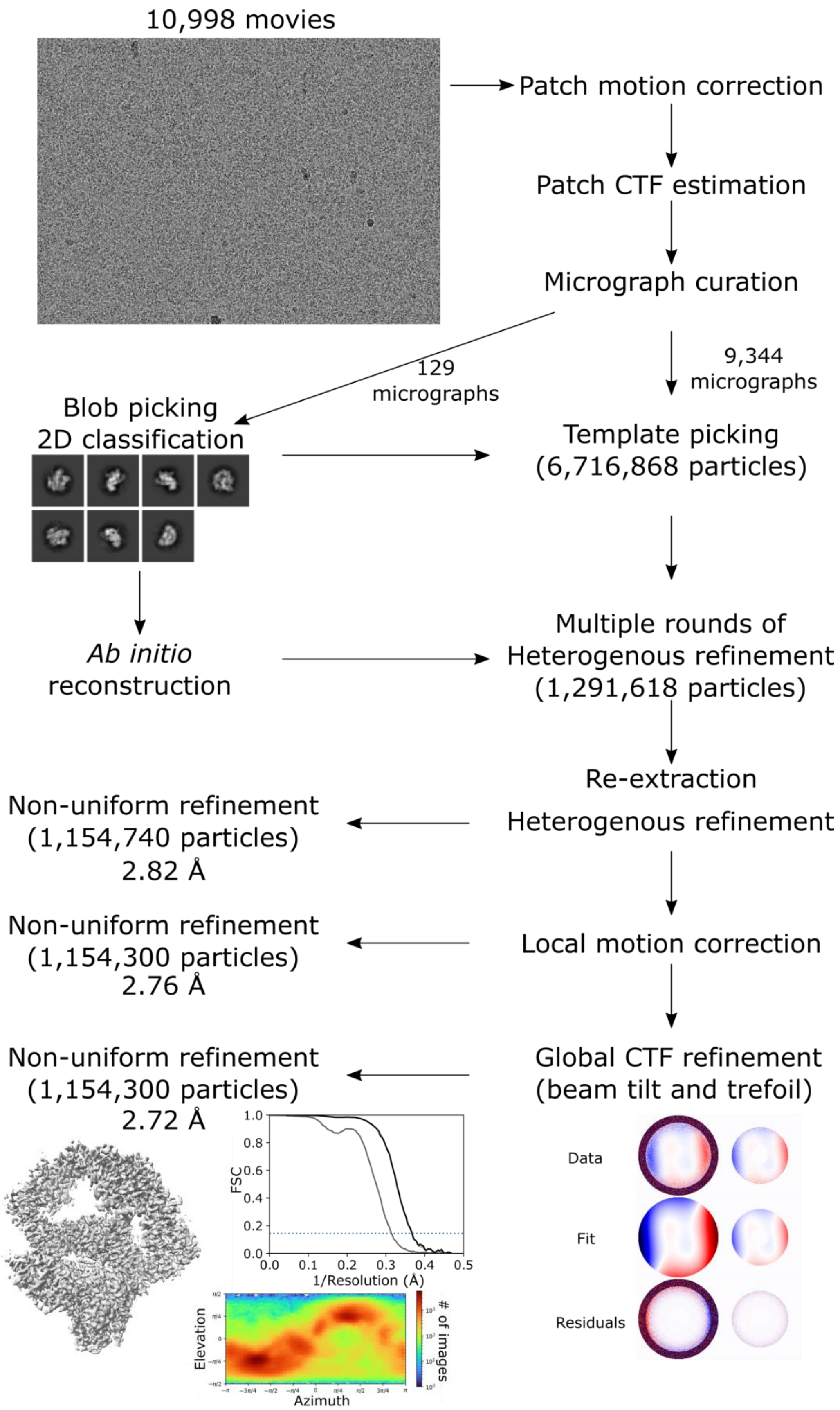


**Fig. S1.** Cryo-EM data particle processing workflow to yield a consensus reconstruction for USP1-UAF1-FANCI-FANCD2$^{Ub}$-ML323 structure. Half-map FSCs for no mask (dark gray), a tight mask (light gray) and corrected (black) and distributions of particle orientations are shown for the final reconstruction.

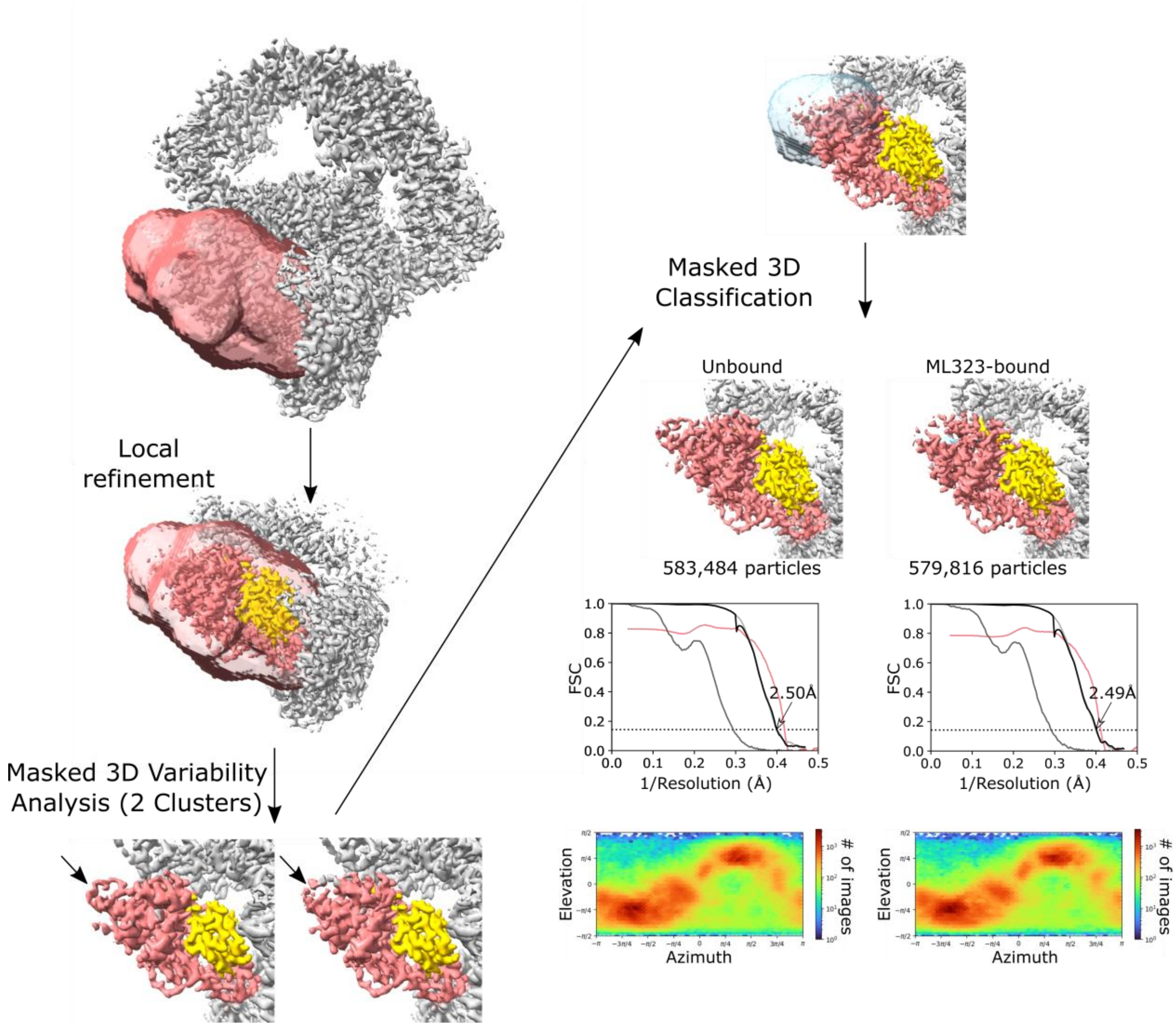


**Fig. S2.** Local refinement and classification of the USP1-ubiquitin subcomplex. Local refinement using a mask around USP1 and ubiquitin was performed, followed by 3D variability analysis solving for 2 modes and using the same mask. Particles were clustered into two groups revealing heterogeneity at the tip of USP1 indicated by arrows. Maps from these clusters were used as inputs for 3D classification with a mask around the surrounding region yielding two subsets of particles – one with density consistent with ML323 and one without. Half-map FSCs for no mask (dark gray), a tight mask (light gray) and corrected (black), as well as the model-map FSC (pink) are shown for each subset. Distributions of particle orientations are shown for each subset.

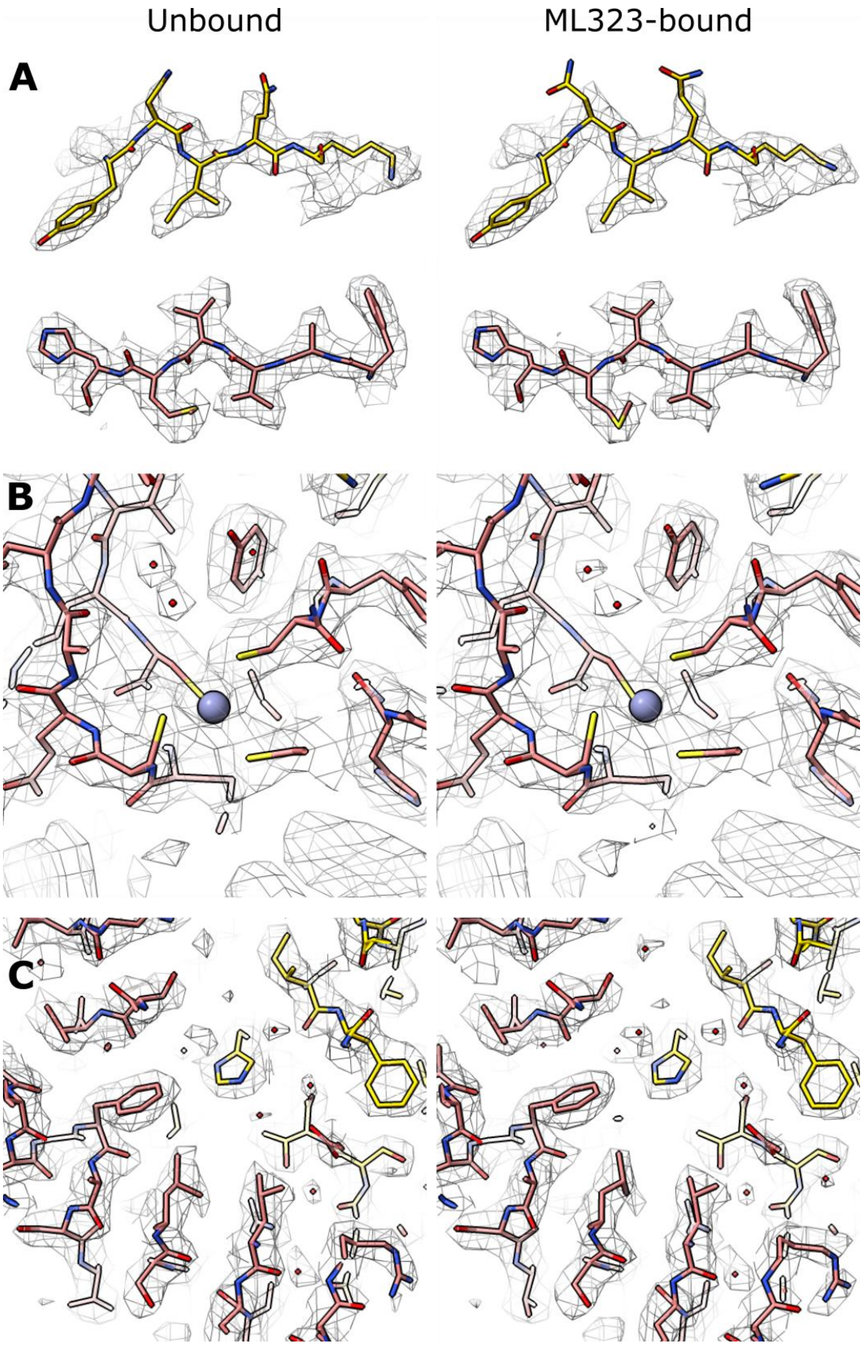


**Fig. S3.** Example density from cryo-EM maps with the fitted models. (**A**) β-strands from ubiquitin and USP1. (**B**) The USP1 zinc fingers. (**C**) The USP1-ubiquitin interface.

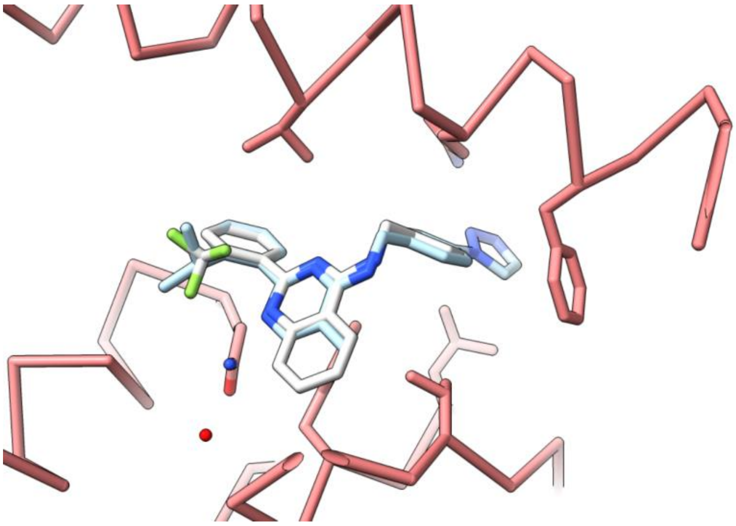

**Fig. S4.** Modeling of the initial hit (white) from which ML323 (blue) was derived (*10*), based on the cryo-EM map and ML323 structure.

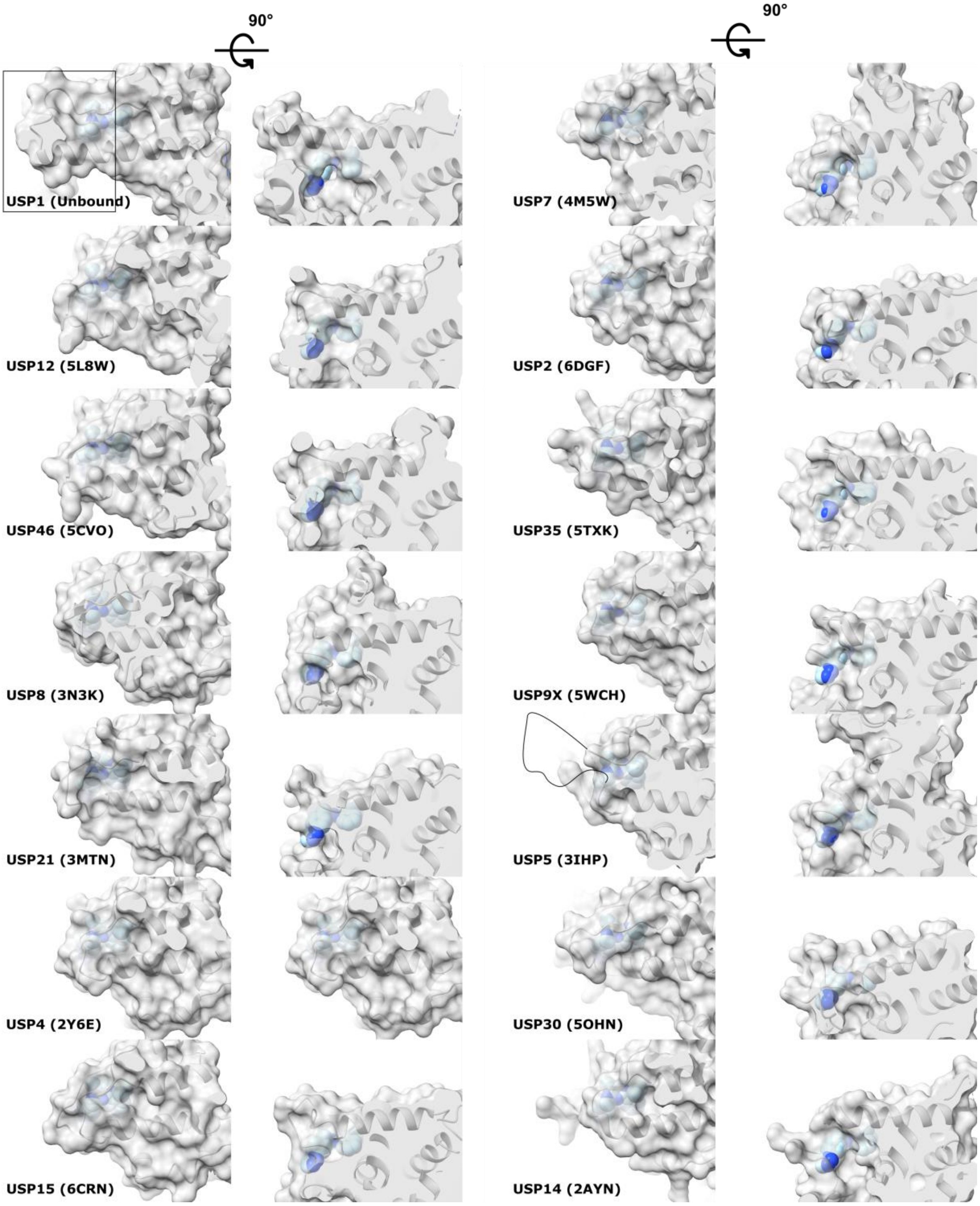


**Fig. S5.** Overlay of the ML323 binding site onto other USP structures *(22, 23, 42–50)*. Only USP1 has an extended helix in the boxed region adjacent to the ML323 binding site. USP5 has an insert in this region however in the 90° rotation, where there is a pocket in USP1, there is none in USP5.

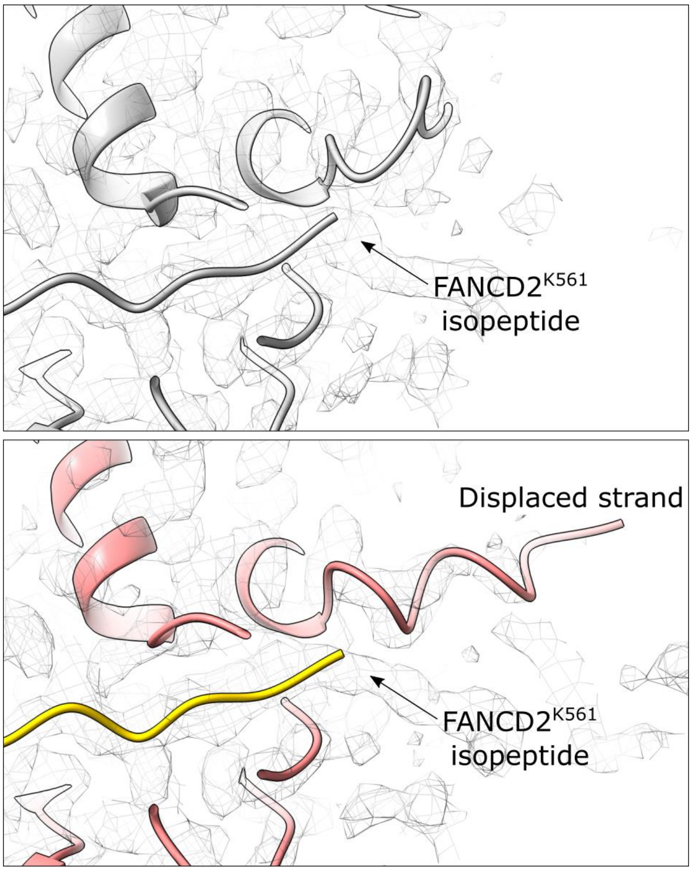


**Fig. S6.** Potential steric hinderance between USP1 and substrate in the ML323-bound state. The unbound state (top) and ML323-bound state (bottom) are compared around the isopeptide bond of the substrate.

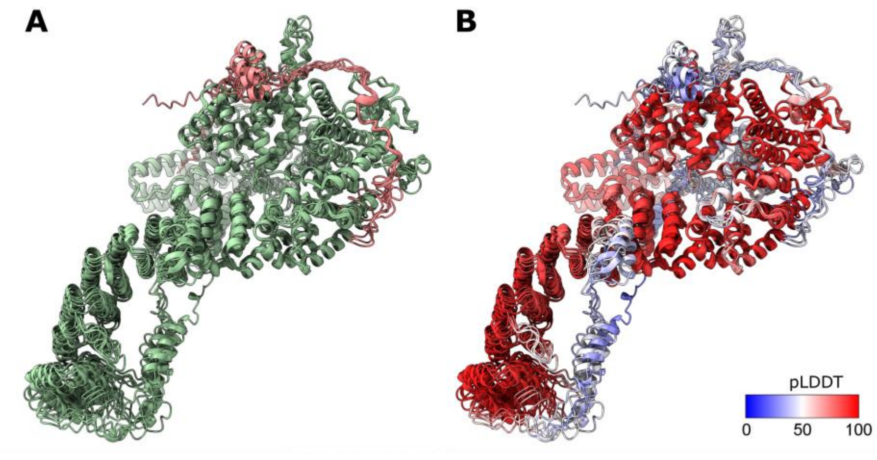


**Fig. S7.** (**A**) Five AlphaFold models of FANCD2 (green) and the USP1 N-terminal extension (NTE) (pink). (**B**) Five AlphaFold models of FANCD2 and the USP1 NTE, all colored by pLDDT confidence score. For USP1, only the region of relatively high confidence (residues 21-25) superimposes well.

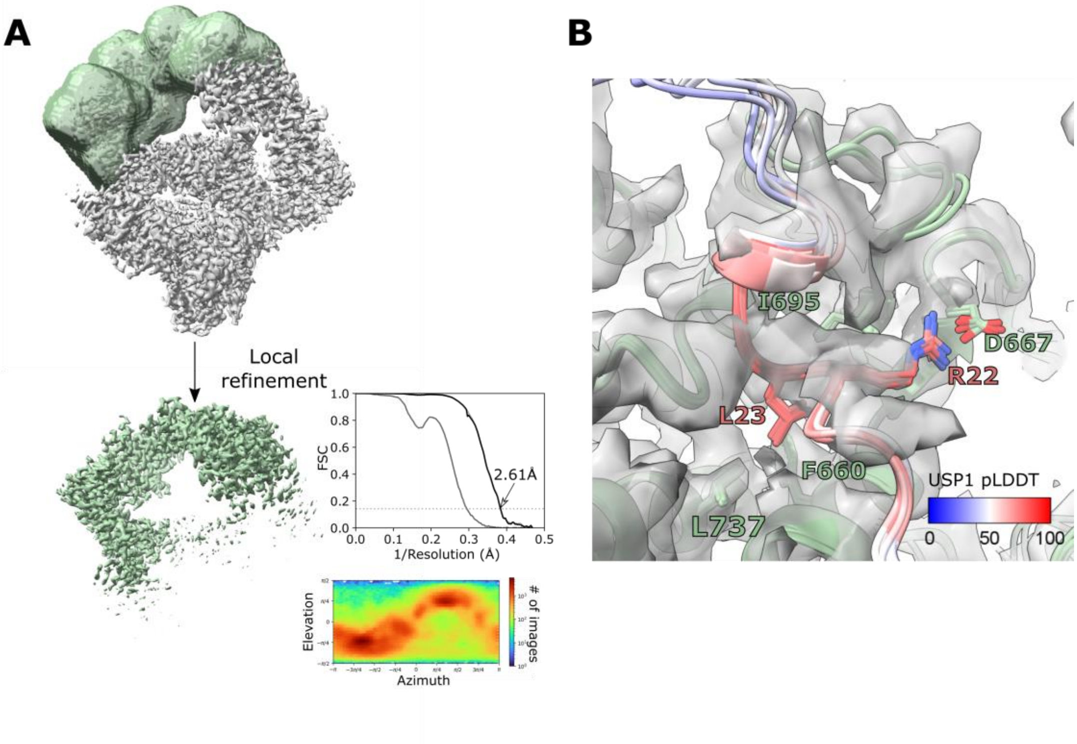


**Fig. S8.** (**A**) Local refinement of the FANCD2 helical domain and C-terminal domain. Half-map FSCs for no mask (dark gray), a tight mask (light gray) and corrected (black) are shown. Distribution of particle orientations is shown. (**B**) Five AlphaFold models fitted into the cryo-EM map locally refined around the FANCD2 C-terminal domain. The USP1 NTE is colored by pLDDT confidence score. Key sidechains involved in the interaction are shown.

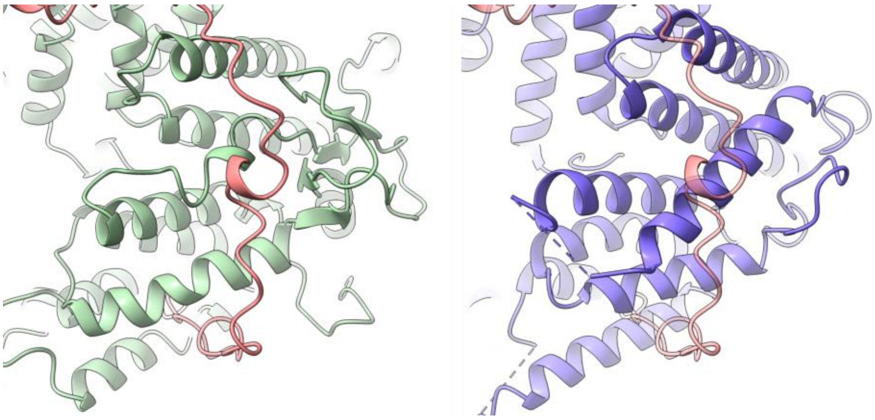

**Fig. S9.** Comparison of the NTE binding site on FANCD2 with the equivalent region on FANCI (6VAD) (*17*). FANCI and FANCD2 were aligned by residues 600-800.

**Table S1.** Cryo-EM data collection and model refinement statistics

| | Consensus | USP1-ubiquitin | USP1-ubiquitin-ML323 | FANCD2$^{CTD}$ |
|---|---|---|---|---|
| **Data collection and processing** | | | | |
| Microscope | Krios | | | |
| Detector | K3 | | | |
| Nominal Magnification | 81,000x | | | |
| Voltage (kV) | 300 | | | |
| Electron Dose ($e^-/Å^2$) | 40 | | | |
| Defocus range (μm) | -1.0 to -3.0 | | | |
| Pixel Size (Å) | 1.06 | | | |
| Symmetry imposed | C1 | | | |
| Map resolution (Å) | 2.72 | 2.50 | 2.49 | 2.61 |
| FSC threshold | 0.143 | 0.143 | 0.143 | 0.143 |
| Map resolution range (Å)[a] | 2.3-6.3 | 2.3-5.5 | 2.3-5.5 | 2.3-4.0 |
| FSC threshold | 0.143 | 0.143 | 0.143 | 0.143 |
| EMDB ID | EMD-XXXXX | EMD-XXXXX | EMD-XXXXX | EMD-XXXXX |
| **Refinement** | | | | |
| Initial models used | | 7AY1 (chains C and D) | 7AY1 (chains C and D) | |
| Map sharpening B-factor ($Å^2$) | | 81.5 | 80.8 | |
| Correlation coefficient (mask)[b] | | 0.86 | 0.83 | |
| Bond length rmsd (Å) | | 0.010 | 0.004 | |
| Bond angle rmsd (°) | | 0.842 | 0.623 | |
| All-atom clashscore | | 6.87 | 4.63 | |
| Ramachandran plot | | 0.00 | 0.00 | |
| Outliers (%) | | 3.82 | 2.31 | |
| Favored (%) | | 96.18 | 97.69 | |
| Rotamer outliers (%) | | 0.00 | 0.00 | |
| PDB ID | | XXXX | XXXX | |

[a]1% and 99% quantiles

[b]Calculated in phenix

**Table S2.**

| | Purification step (column)/Experiment | Buffer composition |
|---|---|---|
| USP1 or UAF1 | Lysis | 50 mM Tris pH 8.0, 150 mM NaCl, 5% glycerol, 10 mM β-mercaptoethanol, 10 mM Imidazole, 2 mM $MgCl_2$, 1x cOmplete EDTA-free protease inhibitor cocktail, >10 units/mL benzonase |
| | Ni-NTA Wash 1/Subtractive | 50 mM Tris pH 8.0, 500 mM NaCl, 5% glycerol, 10 mM β-mercaptoethanol, 10 mM Imidazole |
| | Ni-NTA Wash 2 | 50 mM Tris pH 8.0, 100 mM NaCl, 5% glycerol, 10 mM 1 mM TCEP, 10 mM Imidazole |
| | Ni-NTA Elution | 50 mM Tris pH 8.0, 75 mM NaCl, 5% glycerol, 1 mM TCEP, 250 mM Imidazole |
| | Anion Exchange (ResourceQ 1 mL) | 50 mM Tris pH 8.0, 100-1000 mM NaCl, 5% glycerol, 1 mM TCEP |
| | Gel Filtration (Superdex 200 Increase 10/300 GL) | 20 mM Tris pH 8.0, 150 mM NaCl, 5% glycerol, 5 mM DTT |
| FANCD2, FANCI | Lysis | 50 mM Tris pH 8.0, 400 mM NaCl, 5% glycerol, 5 mM β-mercaptoethanol, 10 mM Imidazole, 2 mM $MgCl_2$, 1x cOmplete EDTA-free protease inhibitor cocktail, >10 units/mL benzonase |
| | Ni-NTA Wash 1 | 50 mM Tris pH 8.0, 400 mM NaCl, 5% glycerol, 5 mM β-mercaptoethanol, 10 mM Imidazole |
| | Ni-NTA Wash 2 | 50 mM Tris pH 8.0, 150 mM NaCl, 5% glycerol, 1 mM TCEP, 10 mM Imidazole |
| | Ni-NTA Elution | 50 mM Tris pH 8.0, 100 mM NaCl, 5% glycerol, 1 mM TCEP, 250 mM Imidazole |
| | Anion Exchange (HP Q 5 mL) | 50 mM Tris pH 8.0, 100-1000 mM NaCl, 5% glycerol, 1 mM TCEP |
| | Gel Filtration (Superose 6 Increase 10/300 GL) | 20 mM Tris pH 8.0, 400 mM NaCl, 5% glycerol, 5 mM DTT |
